# Genetic factors underlying the bidirectional relationship between autoimmune and mental disorders – findings from a Danish population-based study

**DOI:** 10.1101/699462

**Authors:** Xueping Liu, Ron Nudel, Wesley K. Thompson, Vivek Appadurai, Andrew J. Schork, Alfonso Buil, Simon Rasmussen, Rosa L. Allesøe, Thomas Werge, Ole Mors, Anders D. Børglum, David M. Hougaard, Preben B. Mortensen, Merete Nordentoft, Michael E. Benros

**Affiliations:** CORE-Copenhagen Research Centre for Mental Health, Mental Health Centre Copenhagen, Copenhagen University Hospital, Copenhagen, Denmark; iPSYCH, The Lundbeck Foundation Initiative for Integrative Psychiatric Research, Denmark; Institute of Biological Psychiatry, Mental Health Centre Sct. Hans, Mental Health Services Copenhagen, Roskilde, Denmark; Department of Family Medicine and Public Health, Division of Biostatistics, University of California, San Diego, California, USA; Novo Nordisk Foundation Center for Protein Research, Faculty of Health and Medical Sciences, University of Copenhagen, Copenhagen, Denmark; Department of Clinical Medicine, Faculty of Health and Medical Sciences, University of Copenhagen, Copenhagen, Denmark; Psychosis Research Unit, Aarhus University Hospital, Risskov, Denmark; Department of Biomedicine, Aarhus University and Centre for Integrative Sequencing, iSEQ, Aarhus, Denmark; Aarhus Genome Center, Aarhus, Denmark; Center for Neonatal Screening, Department for Congenital Disorders, Statens Serum Institut, Copenhagen, Denmark; National Center for Register-Based Research, Aarhus University, Aarhus, Denmark

**Keywords:** autoimmune diseases, mental disorders, comorbidity, bidirectionality, human leukocyte antigen, genome-wide association study

## Abstract

**Background:** Previous studies have indicated the bidirectionality between autoimmune and mental disorders. However, genetic studies underpinning the co-occurrence of the two disorders have been lacking. In this study, we examined the potential genetic contribution to the association between autoimmune and mental disorders.

**Methods:** We used diagnostic information for patients with seven autoimmune diseases and six mental disorders from the Danish population-based case-cohort sample (iPSYCH2012). We explored the epidemiological association using survival analysis and modelled the effect of polygenic risk scores (PRSs) on two diseases. The genetic factors were investigated using GWAS and HLA imputation data based on iPSYCH cohort.

**Results:** Among 64,039 individuals, a total of 43,902 (68.6%) were diagnosed with mental disorders and 1,383 (2.2%) with autoimmune diseases. There was a significant comorbidity between the two diseases (*P*=2.67×10^-7^, OR=1.38, 95%CI=1.22-1.56), with an overall bidirectional association wherein individuals with autoimmune diseases had an increased risk of subsequent mental disorders (HR=1.13, 95%CI: 1.07-1.21, *P*=7.95×10^-5^) and *vice versa* (HR=1.27, 95%CI=1.16-1.39, *P*=8.77×10^-15^). Though PRSs were significantly correlated with both types of diagnosis, PRSs had little effect on the bidirectional relationship. Importantly, we for the first time observed 12 human leukocyte antigen (HLA) loci and 20 HLA alleles strongly associated with overall autoimmune diseases, but we did not find significant evidence of their associations with overall mental disorders.

**Conclusions:** Our findings confirm the overall comorbidity and bidirectionality between autoimmune and mental disorders and discover HLA genes which are significantly associated with overall autoimmune diseases, but not with overall mental disorders.

## Introduction

Individuals with autoimmune diseases have an increased risk of mental disorders, such as schizophrenia (1, 2), depression (3, 4) and bipolar disorder (5). Studies have also shown that individuals with mental disorders such as schizophrenia (6, 7) and depression (3, 4, 8) have a subsequently increased risk of autoimmune diseases. Furthermore, this association between autoimmune diseases and mental disorders has been confirmed in a recent meta-analysis (9). However, not all autoimmune diseases are associated with mental disorders (9), and there is a well-documented negative association between rheumatoid arthritis and schizophrenia (10).

Given the overall substantial genetic component to both disease classes, it has been suggested that some of the epidemiologically observed co-occurrence of autoimmune and mental disorders could be explained by shared genetic risk factors. Several studies have attempted to estimate the genetic correlation between them; however, these have resulted in conflicting effects (positive, negative or even no correlation). Pouget *et al.* (11) examined a comprehensive cross-disorder analysis of schizophrenia with 19 autoimmune and immune diseases using the publicly available genome-wide association study (GWAS) summary statistics and observed small positive genetic correlations of schizophrenia with only a subset of specific immune diseases (r_g_=0.10-0.18), including inflammatory bowel disease, Crohn’s disease, ulcerative colitis, primary biliary cirrhosis, psoriasis and systemic lupus erythematosus. In contrast, a significant negative single nucleotide polymorphism (SNP)-genetic correlation between schizophrenia and rheumatoid arthritis has also been reported (10). Andreassen *et al.* (12) observed that some HLA alleles are significantly enriched in both schizophrenia and multiple sclerosis, but with opposite directions of effect in both diseases; further, they found no genetic overlap between bipolar disorder and multiple sclerosis. Hoeffding *et al.* (13) found no common genetic basis between schizophrenia and autoimmune diseases by connecting significant risk variants of 25 specific autoimmune diseases with schizophrenia. Hence, a larger study may be needed to systematically investigate the overall association between autoimmune and mental disorders.

In this study, we investigated the epidemiological association between the two disease classes using the Danish population-based case-cohort sample (iPSYCH2012) (14), a large, population-based study. To discover the overlapping genes between the two disease classes, we conducted a GWAS for overall autoimmune diseases (i.e. with at least one specific autoimmune disease) and associated the top variants with overall mental disorders (i.e. with at least one specific mental disorder).

## Materials and Methods

### Study population

We investigated all Danes from the large Danish population-based case-cohort sample (iPSYCH2012) from the Integrative Psychiatric Research Consortium (iPSYCH) cohort (14). iPSYCH2012 case-cohort sample consists of 87,764 Danish individuals, who were born in Denmark between 1981 and 2012 with dry blood spots available in Danish Neonatal Screening Biobank (15). In this cohort, there are 57,764 people with at least one major mental disorder and 30,000 heathy controls who were randomly sampled from the population without regard for psychiatric disorders. Genotyping, data cleaning and imputation processes are described in detail by Schork *et al.* (16), resulting in 65,534 subjects with 8,018,013 imputed genetic variants based on integrated phase III release of the 1000 Genomes Project (17). SNP positions were based on National Center for Biotechnology Information (NCBI) build 37 (hg19) and alleles were labelled on the positive strand of the reference genome. In addition to the previous cleaning procedure, we required that the all individuals in the cohort should be born, alive and reside in Denmark by the end of follow-up (31 December 2012), which leads to 64,039 subjects in our study.

### Assessment of mental disorders

People with mental disorders, as diagnosed by the patient’s treating psychiatrist, were included in our study. Date of illness onset was defined as the first day of the first hospital contact for psychiatric diagnosis.

Six mental disorders cases with at least one indication according to the corresponding International Classification of Diseases (ICD, version 8 through 1993 and version 10 from 1994) ICD10 (or equivalent ICD8) codes were identified: schizophrenia (F20), bipolar disorder (F30-31), affective disorders (F32-F39), autism spectrum disorders (F84.0, F84.1, F84.5, F84.8 and F84.9), attention deficit/hyperactivity disorder (ADHD, F90.0), anorexia (F50.0 and F50.1). As an additional case group, we defined those patients with at least one psychiatric indication (overall mental disorders, N=43,902). Controls for overall mental disorders were defined as the individuals without any of these diseases (N=20,137).

### Assessment of autoimmune diseases

The time of onset of an autoimmune disease was defined as the day of first hospitalization with ICD code for one of the diagnosis. Each person could have a history of more than one autoimmune disease. We omitted those autoimmune diseases, where the number of patients was lower than 50. In total, we obtained seven specific autoimmune diseases, including Crohn’s disease (ICD8: 56301, ICD10: K50), ulcerative colitis (ICD8: 56319, ICD10: K51), celiac diseases (ICD8: 26900, ICD10: K900), rheumatoid arthritis (ICD8: 71219, 71239 and 71259, ICD10: M05 and M06), psoriasis (ICD8: 69609, 69610 and 69619, ICD10: L400-L403, L405-L409), type 1 diabetes (ICD8: 249, ICD10: E10) and juvenile idiopathic arthritis (ICD8: 71209, ICD10: M08). Similar to the assessment of mental disorders, we defined an additional group of patients as having at least one of these seven diseases (overall autoimmune diseases, N=1,383). Controls for overall autoimmune diseases were defined as those individuals without any autoimmune diseases (N=62,656).

### Selection criteria for public GWAS summary statistics

To not include the bias for polygenic risk scoring, we required that the publicly GWAS results that were most recently published for autoimmune and mental disorders, should be conducted on non-Danish samples.

### PRSs for autoimmune and mental disorders

Here we used the software package PRSice-2 (18) and its default parameter settings to calculate PRSs for each individual based on the GWAS summary statistics. Specifically, if any pairs of SNPs in the GWAS results have a physical distance smaller than 250kb, and *r*^*2*^ higher than 0.1, the less significant SNP is removed. Afterwards, we calculated the PRSs as the sum of the remaining SNPs weighted by their marginal effect estimates. These SNPs are largely independent with GWAS *P* values below a threshold *P*_*T*_. In our study, we considered *P*_*T*_ ∈ {1E-8, 1E-7, 1E-6, 1E-5, 3E-5, 1E-4, 3E-4, 0.001, 0.003, 0.01, 0.03, 0.1, 0.3, 1} and reported the best fit across these thresholds calculated by PRSice software. Both HLA and non-HLA regions were included to generate PRSs for autoimmune diseases. The PRSs for overall autoimmune diseases is the sum of all the scores of seven autoimmune PRSs. Similarly, PRSs for six specific and overall mental disorders were calculated.

### GWAS analysis

We conducted the GWAS scan for having overall autoimmune diseases using SNPTEST (19). Controls were defined as subjects without any of the seven autoimmune diseases. Association testing was run using logistic regression based on imputed allelic dosage data under an additive genetic model, adjusting for mental disorder status (overall mental disorders) and the first four components from the principal components’ analysis.

### Identification of independent loci

For the genome-wide significant SNPs (*P*< 5×10^-8^) from overall autoimmune diseases, we generated independent “lead SNPs” by keeping the strongest evidence of being associated from each set of SNPs that are in linkage disequilibrium (LD pairwise r^2^>0.05) and/or in proximity (physical distance<1Mb). These “lead SNPs” were defined as our independent loci. This procedure was performed using PLINK 1.9 (20) (www.cog-genomics.org/plink/1.9/): --clump-p1 5e-8 -- clump-kb 500 --clump-r2 0.05. The imputed data of iPSYCH2012 cohort was used to calculate LD between the SNPs that feature in the GWAS results.

### Associations of HLA alleles with autoimmune and mental disorders

The gene-based tests and the allele-specific tests were performed using the procedure described in Nudel *et al.* (21), with the same covariates described there. The tests for the overall autoimmune diseases were run with an additional covariate for having an ICD10 chapter V diagnosis from the psychiatric register. Gene-based tests significant after false discovery rate (FDR) (22) correction (q ≤ 0.05) were followed up with allele-specific tests for the relevant phenotype-gene associations.

### Statistical analysis

The chi-squared test was applied to examine the epidemiological correlation between autoimmune and mental disorders. Odds ratio (OR) and 95% confidence intervals (95%CI) were calculated using the “epitools” package in R (23).

We followed the method proposed by Euesden *et al.* (4) to investigate the time-course of overall mental disorders on overall autoimmune disease onset and *vice versa*. Namely, we firstly generated multiple observations per individual: one before exposure to the outcome (autoimmune / mental), one before the predictor (mental / autoimmune), and a third when the individual is exposed to both diseases until the most recent point when the data was collected (it is 31 December 2012 for the iPSYCH2012 cohort). Afterwards, Breslow’s method was applied to estimate the baseline hazard function using Cox Proportional Hazards (CoxPH) model, adjusted for sex and four principal components.

All PRSs for different autoimmune and mental disorders were standardized mean=0 and SD=1. Logistic regression was applied to examine whether PRSs for autoimmune diseases predict the diagnosis of autoimmune diseases, and whether PRSs for mental disorders predict the mental disorders status. All the statistical analyses were performed using R (23).

## Results

### Sample characteristics

We restricted our analyses into the 64,039 individuals in the iPSYCH2012 cohort which survived after a series of data cleaning processes (see Materials and Methods). There were 43,902 individuals with overall mental disorders, including schizophrenia (N=2,117), bipolar disorder (N=1,185), affective disorder (N=19,146), autism spectrum disorders (N=11,933), attention deficit/hyperactivity disorder (ADHD, N=13,955), and anorexia (N=2,772). Similarly, we identified 1,383 patients with overall autoimmune diseases, encompassing seven specific diseases, namely, Crohn’s disease (N=227), ulcerative colitis (N=280), celiac diseases (N=122), rheumatoid arthritis (N=97), psoriasis (N=143), type 1 diabetes (N=424) and juvenile idiopathic arthritis (N=237). The average age at onset for overall autoimmune disease was 20.11 (SD=6.42), and for mental disorders the mean age was 16.87 (SD=6.71). Among the 43,902 individuals diagnosed with any mental disorders, a total of 1,036 were also diagnosed with an autoimmune disease (2.36%, **Table 1**). Among the 20,137 individuals without mental disorders, a total of 347 (1.72%) were diagnosed with autoimmune diseases. There is a significant correlation between having overall autoimmune and mental disorders (Chi-square test, *P*=2.67× 10^-7^, OR=1.38, 95% CI=1.22-1.56), indicating the two diseases co-occur in the same individuals at a greater rate than would be expected by chance.

**Table 1:** Sample characteristics (born between 1981 and 2012)

| Disease status | #Total | Crohn's disease | Ulcerative colitis | Celiac diseases | Rheumatoid arthritis | Psoriasis | Type 1 diabetes | Juvenile idiopathic arthritis | Overall autoimmune diseases |
| --- | --- | --- | --- | --- | --- | --- | --- | --- | --- |
| Overall mental disorders | 43,902 | 169 | 210 | 96 | 83 | 104 | 324 | 159 | 1,036 |
| Schizophrenia | 2,117 | 11 | 13 | 5 | 4 | 9 | 24 | 12 | 70 |
| Bipolar disorder | 1,185 | 6 | 6 | <3 | <3 | 4 | 13 | <3 | 28 |
| Affective disorder | 19,146 | 119 | 146 | 47 | 60 | 60 | 172 | 78 | 616 |
| Autism spectrum disorders | 1,1933 | 21 | 28 | 29 | 6 | 13 | 81 | 40 | 199 |
| ADHD | 13,955 | 32 | 27 | 19 | 8 | 27 | 68 | 37 | 204 |
| Anorexia | 2,772 | 18 | 23 | 8 | 13 | 13 | 26 | 17 | 101 |
| Controls | 20,137 | 58 | 70 | 26 | 14 | 39 | 100 | 78 | 347 |

### Bidirectionality between autoimmune and mental disorders

Next, we evaluated the bidirectional relationship of the two disease groups. For that, we applied CoxPH models to evaluate whether there is an increased risk of mental disorders in individuals with autoimmune diseases, and *vice versa*, adjusting for sex and four principal components. We found that a hospital contact with overall autoimmune disease was significantly associated with an increased risk of overall mental disorders (hazard ratio HR=1.13, 95%CI: 1.07-1.21, *P*=7.95×10^-5^) (**Table 2**). We also found significant evidence for an effect of mental disorder onset increasing hazard of subsequent overall autoimmune diagnosis (HR=1.27, 95%CI=1.16-1.39, *P*=8.77×10^-15^). These results point an overall bidirectional relationship between the two disease groups. However, when we investigated the specific associations between them (i.e. specific autoimmune disease *versus* overall mental disorders, specific mental disorder *versus* overall autoimmune disease), the results vary from no (e.g. overall mental disorders *versus* psoriasis), negative (e.g. overall mental disorders *versus* ulcerative colitis) to positive associations (**Table S1**), evidencing the complex association between the two types of illness.

**Table 2:** Bidirectional association between autoimmune and mental disorders (born between 1981and 2012)

|  | PRS for overall autoimmune diseases |  | Basic adjustment * |  | Additional adjustment for autoimmune PRS |  | Additional adjustment for mental PRS |  | Additional adjustment for autoimmune PRS and mental PRS |  |
| --- | --- | --- | --- | --- | --- | --- | --- | --- | --- | --- |
|  | *P* | HR (95% CI) | *P* | HR (95% CI) | *P* | HR (95% CI) | *P* | HR (95% CI) | *P* | HR (95% CI) |
| Overall mental disorders | 0.038 | 1.01 (1.00 – 1.02) | 7.95E-5 | 1.13 (1.07 – 1.21) | 6.97E-5 | 1.14 (1.07 – 1.21) | 1.03E-4 | 1.13 (1.06 – 1.2) | 8.61E-5 | 1.13 (1.07 – 1.21) |

B. Effect of having mental disorders on autoimmune diseases
|  | PRS for overall mental disorders |  | Basic adjustment * |  | Additional adjustment for autoimmune PRS |  | Additional adjustment for mental PRS |  | Additional adjustment for autoimmune PRS and mental PRS |  |
| --- | --- | --- | --- | --- | --- | --- | --- | --- | --- | --- |
|  | *P* | HR (95% CI) | *P* | HR (95% CI)<sup>a</sup> | *P* | HR (95% CI) | *P* | HR (95% CI) | *P* | HR (95% CI) |
| Overall autoimmune diseases | 0.42 | 1.02 (0.98 – 1.07) | 8.77E-15 | 1.27 (1.16 – 1.39) | 1.95E-14 | 1.27 (1.16 – 1.39) | 1.63E-14 | 1.27 (1.6 – 1.39) | 2.96E-14 | 1.27 (1.16 – 1.38) |

\*, adjusted for sex and four principal components.

### Effect of PRSs on the bidirectionality

We retrieved public GWAS summary statistics for autoimmune and mental disorders and calculated PRSs for each specific and overall autoimmune/mental diseases in our cohort study using various GWAS *P* value thresholds (see **Materials and Methods**). Afterwards, we performed a logistic regression analysis using the PRSs as the predictors of disease onset. We found that all PRSs for autoimmune diseases were significantly correlated with the autoimmune diagnosis with an increased odds ratio (OR), same scenario for mental PRSs associated with mental disorders (lowest OR value is: 1.09, with 95%CI: 1.07-1.11, **Table 3**).

**Table 3:** Summary of public GWAS data and associations between PRS and diseases

|  | Public GWAS statistics | PRS calculations |
| --- | --- | --- |

| Autoimmune / Mental disorders | #cases | #controls | #snps<sup>a</sup> | PMID | #snps<sup>b</sup> | OR (95%CI)<sup>c</sup> | P<sup>c</sup> |
| --- | --- | --- | --- | --- | --- | --- | --- |
| Overall mental disorders |  |  |  |  |  | 1.12 (1.1,1.14) | 1.98E-38 |
| Schizophrenia | 35,476 | 46,839 | 9,444,230 | 25056061 | 135,904 | 1.28 (1.22, 1.34) | 1.51E-25 |
| Bipolar disorder | 7,647 | 27,303 | 10,082,186 | 27329760 | 22,409 | 1.24 (1.17,1.32) | 1.43E-12 |
| Affective disorder | 9,240 | 9,519 | 1,235,109 | 22472876 | 38,500 | 1.09 (1.07,1.11) | 1.23E-16 |
| Autism spectrum disorders | 7,387 | 8,567 | 6,440,259 | 28540026 | 49,007 | 1.15 (1.12,1.18) | 8.56E-31 |
| ADHD | 896 | 2,455 | 1,206,461 | 20732625 | 37,538 | 1.12 (1.1,1.15) | 1.49E-24 |
| Anorexia | 3,495 | 10,982 | 10,641,224 | 28494655 | 56,890 | 1.22 (1.17,1.27) | 3.17E-20 |
| Overall autoimmune diseases |  |  |  |  |  | 1.24 (1.17,1.31) | 6.03E-14 |
| Crohn's disease | 12,194 | 28,072 | 9,570,787 | 28067908 | 338 | 1.57 (1.38,1.8) | 2.09E-11 |
| Ulcerative colitis | 12,366 | 33,609 | 9,588,016 | 28067908 | 1,127 | 1.37 (1.22,1.54) | 1.19E-7 |
| Celiac diseases | 11,812 | 11,837 | 131,579 | 22057235 | 18,628 | 1.86 (1.48,2.32) | 6.79E-8 |
| Rheumatoid arthritis | 19,234 | 43,923 | 9,739,303 | 24390342 | 2,243 | 1.39 (1.15,1.67) | 5.82E-4 |
| Psoriasis | 2,997 | 9,183 | 161,173 | 23143594 | 2,354 | 1.87 (1.64,2.13) | 1.9E-20 |
| Type 1 diabetes | 6,683 | 12,173 | 121,694 | 25751624 | 566 | 1.74 (1.59,1.92) | 1.33E-28 |
| Juvenile idiopathic arthritis | 2,816 | 13,056 | 122,330 | 23603761 | 169 | 1.37 (1.2,1.57) | 3.55E-6 |

a, number of snps in the public GWAS summary statistics.
b, number of snps included in PRSs calculations.
c, logistic models were adjusted for sex and four principal components.

Next, we examined whether PRSs for overall mental disorders predict overall autoimmune diseases diagnosis, and whether PRSs for overall autoimmune diseases predict overall mental disorders diagnosis using the CoxPH models. We observed that PRSs for overall mental disorders were not associated with overall autoimmune diseases onset. Though overall autoimmune PRSs were associated with the overall mental disorders (*P*=0.038, HR=1.01, 95%CI: 1.00-1.02), the highest quintile of PRSs was not significantly associated with an increased hazard of overall mental disorders, with CI including the null value (*P*=0.062, HR=1.04, 95%CI: 0.998-1.09, **Figure S1**). To examine the effect of PRSs on the bidirectional association, we used overall PRSs as the covariates in the previous bidirectional models, and we observed that additional adjustment for autoimmune, mental disorders and both overall PRSs did not alter the general associations between autoimmune and mental disorders (**Table 2**). These results confirm that the overall genetic susceptibility has little effect on the bidirectional relationship between the two disease classes.

### Shared genetic risk

The occurrence of overall autoimmune diseases patients with overall mental disorders could reflect genetically determined factors. To investigate the genetic contribution to the bidirectional relationship, we performed a GWAS for overall autoimmune diseases based on the 1,383 patients and 62,656 controls (**Figure 1**). After imputation using reference data from the integrated phase III release of the 1000 Genomes Project, we analyzed 7,721,839 genetic variants with high imputation quality (info score > 0.8, minor allele frequency>0.01). The genomic inflation factor was 1.008, indicating little evidence of population stratification. Our results reveal that genes located in the major histocompatibility complex (MHC) and, in particular loci from human leucocyte antigen (HLA) region, which encode molecules involved in antigen presentation, inflammation, the complement system and the immune responses (24), are strongly associated with overall autoimmune diseases. This region includes a total of 12 independent loci, harboring *HLA-DQB1, HLA-DQA2, AL671883.1, MICA, HLA-B, PSMB9, TAP1, HLA-DOA* and *HLA-DMB, etc*. genes, with rs1064173-A as the SNP most significantly associated with the overall autoimmune diseases (*P*=2.74□× 10^-36^, OR=1.80, 95% CI: 1.64-1.96) (see **Table 4**).

**Figure 1:**
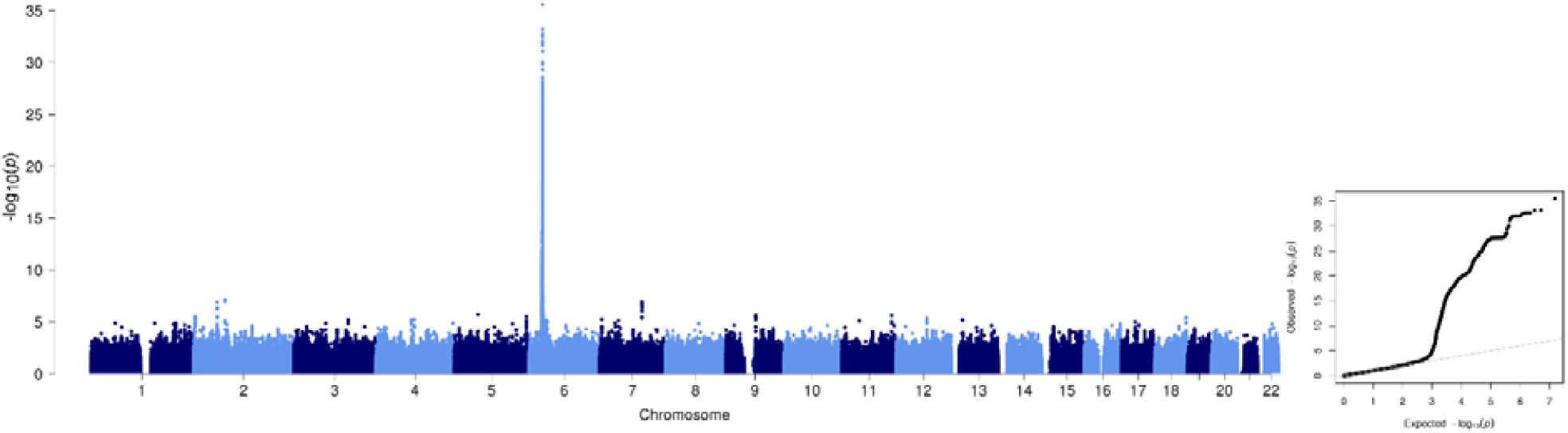
Manhattan plot for having overall autoimmune diseases of -log_10_(*P*) values across the chromosomes (left panel) and corresponding quantile-quantile plot of observed *versus* expected – log_10_(*P*) values (right panel).

**Table 4:**
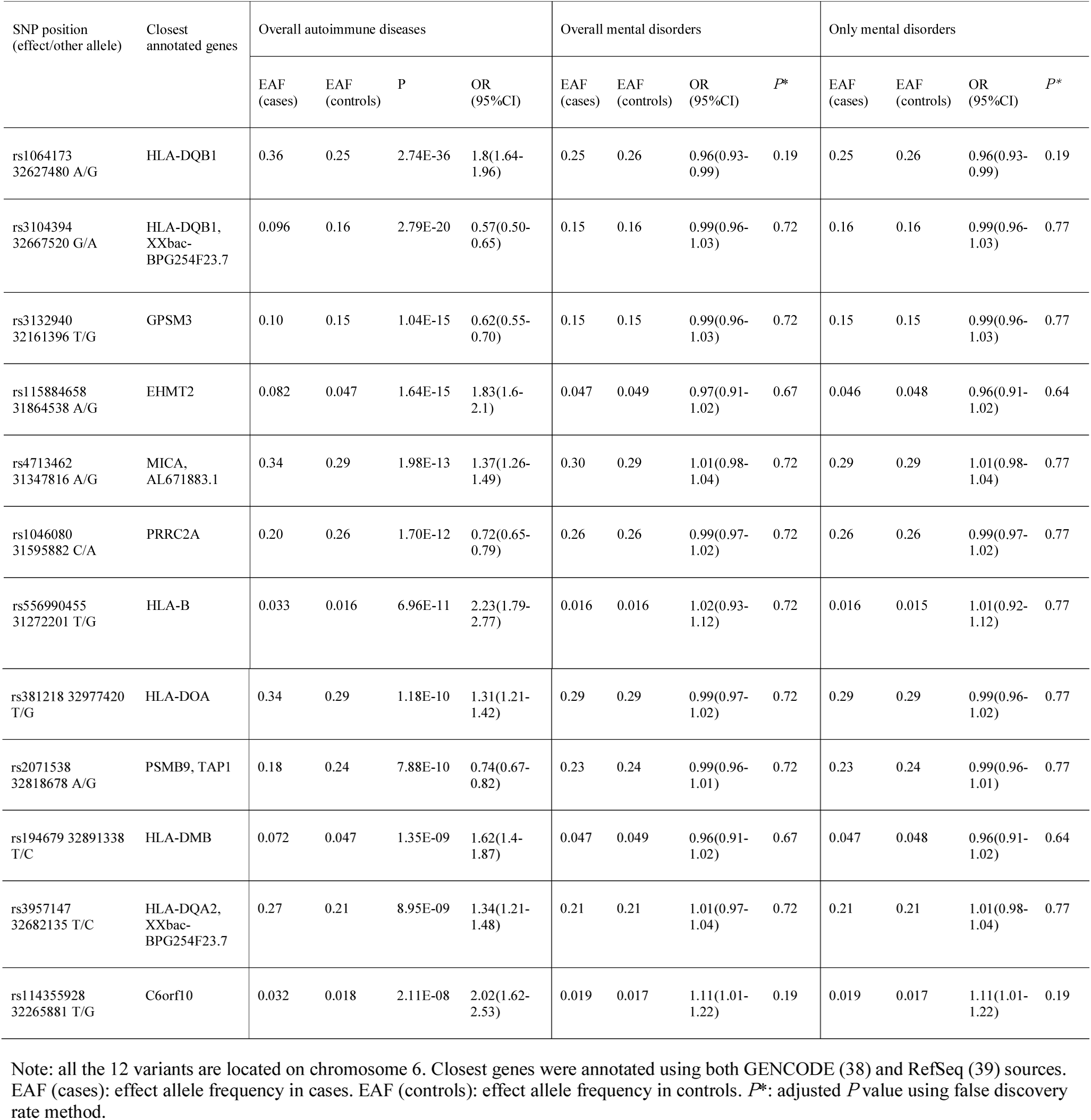
Associations of the autoimmune top variants with mental disorders

| SNP position<br>(effect/other allele) | Closest<br>annotated genes | Overall autoimmune diseases |  |  |  | Overall mental disorders |  |  |  | Only mental disorders |  |  |  |
| --- | --- | --- | --- | --- | --- | --- | --- | --- | --- | --- | --- | --- | --- |
|  |  | EAF<br>(cases) | EAF<br>(controls) | P | OR<br>(95%CI) | EAF<br>(cases) | EAF<br>(controls) | OR<br>(95%CI) | P* | EAF<br>(cases) | EAF<br>(controls) | OR<br>(95%CI) | P* |
| rs1064173<br>32627480 A/G | HLA-DQB1 | 0.36 | 0.25 | 2.74E-36 | 1.8(1.64-1.96) | 0.25 | 0.26 | 0.96(0.93-0.99) | 0.19 | 0.25 | 0.26 | 0.96(0.93-0.99) | 0.19 |
| rs3104394<br>32667520 G/A | HLA-DQB1,<br>XXbac-BPG254F23.7 | 0.096 | 0.16 | 2.79E-20 | 0.57(0.50-0.65) | 0.15 | 0.16 | 0.99(0.96-1.03) | 0.72 | 0.16 | 0.16 | 0.99(0.96-1.03) | 0.77 |
| rs3132940<br>32161396 T/G | GPSM3 | 0.10 | 0.15 | 1.04E-15 | 0.62(0.55-0.70) | 0.15 | 0.15 | 0.99(0.96-1.03) | 0.72 | 0.15 | 0.15 | 0.99(0.96-1.03) | 0.77 |
| rs115884658<br>31864538 A/G | EHMT2 | 0.082 | 0.047 | 1.64E-15 | 1.83(1.6-2.1) | 0.047 | 0.049 | 0.97(0.91-1.02) | 0.67 | 0.046 | 0.048 | 0.96(0.91-1.02) | 0.64 |
| rs4713462<br>31347816 A/G | MICA,<br>AL671883.1 | 0.34 | 0.29 | 1.98E-13 | 1.37(1.26-1.49) | 0.30 | 0.29 | 1.01(0.98-1.04) | 0.72 | 0.29 | 0.29 | 1.01(0.98-1.04) | 0.77 |
| rs1046080<br>31595882 C/A | PRRC2A | 0.20 | 0.26 | 1.70E-12 | 0.72(0.65-0.79) | 0.26 | 0.26 | 0.99(0.97-1.02) | 0.72 | 0.26 | 0.26 | 0.99(0.97-1.02) | 0.77 |
| rs556990455<br>31272201 T/G | HLA-B | 0.033 | 0.016 | 6.96E-11 | 2.23(1.79-2.77) | 0.016 | 0.016 | 1.02(0.93-1.12) | 0.72 | 0.016 | 0.015 | 1.01(0.92-1.12) | 0.77 |
| rs381218 32977420 T/G | HLA-DOA | 0.34 | 0.29 | 1.18E-10 | 1.31(1.21-1.42) | 0.29 | 0.29 | 0.99(0.97-1.02) | 0.72 | 0.29 | 0.29 | 0.99(0.96-1.02) | 0.77 |
| rs2071538 32818678 A/G | PSMB9, TAP1 | 0.18 | 0.24 | 7.88E-10 | 0.74(0.67-0.82) | 0.23 | 0.24 | 0.99(0.96-1.01) | 0.72 | 0.23 | 0.24 | 0.99(0.96-1.01) | 0.77 |
| rs194679 32891338 T/C | HLA-DMB | 0.072 | 0.047 | 1.35E-09 | 1.62(1.4-1.87) | 0.047 | 0.049 | 0.96(0.91-1.02) | 0.67 | 0.047 | 0.048 | 0.96(0.91-1.02) | 0.64 |
| rs3957147 32682135 T/C | HLA-DQA2, XXbac-BPG254F23.7 | 0.27 | 0.21 | 8.95E-09 | 1.34(1.21-1.48) | 0.21 | 0.21 | 1.01(0.97-1.04) | 0.72 | 0.21 | 0.21 | 1.01(0.98-1.04) | 0.77 |
| rs114355928 32265881 T/G | C6orf10 | 0.032 | 0.018 | 2.11E-08 | 2.02(1.62-2.53) | 0.019 | 0.017 | 1.11(1.01-1.22) | 0.19 | 0.019 | 0.017 | 1.11(1.01-1.22) | 0.19 |

Note: all the 12 variants are located on chromosome 6. Closest genes were annotated using both GENCODE (38) and RefSeq (39) sources. EAF (cases): effect allele frequency in cases. EAF (controls): effect allele frequency in controls. $P^*$ : adjusted $P$ value using false discovery rate method.

We then tested whether the lead variants of the 12 independent loci are associated with overall mental disorders. We found that none remained associated with overall mental disorders, neither with only mental illness (i.e. after removing the psychiatric patient cases comorbid with autoimmune disease, controls with autoimmune diseases were also deleted) after multiple testing correction using false discovery rate (FDR) method (**Table 4**).

To further understand the genetic architecture in HLA disease association, we performed an association study of overall autoimmune diseases with seven HLA genes (HLA class I loci: A, B and C; HLA class II loci: DPB1, DQA1, DQB1 and DRB1) with imputed HLA genotypes. After multiple testing correction using FDR for the gene-based LRT chi-squared *P* values, we found HLA-C and four HLA class II genes to be significantly associated with overall autoimmune diseases (**Table 5**, methods in details were described by Nudel *et al.* (20)). We performed *post hoc* analyses to explore the associations of autoimmune diseases with the alleles of the five significant genes. Among the 89 tested alleles, 20 remained significant after FDR correction (q ≤ 0.05), but with different directions of effect on the overall autoimmune diseases (see **Table S2**). The gene based tests did not find significant association for the seven genes with neither overall nor only the combined group of mental disorders without autoimmune diseases (**Table 5**), which coincides with our finding using GWAS data.

**Table 5:**
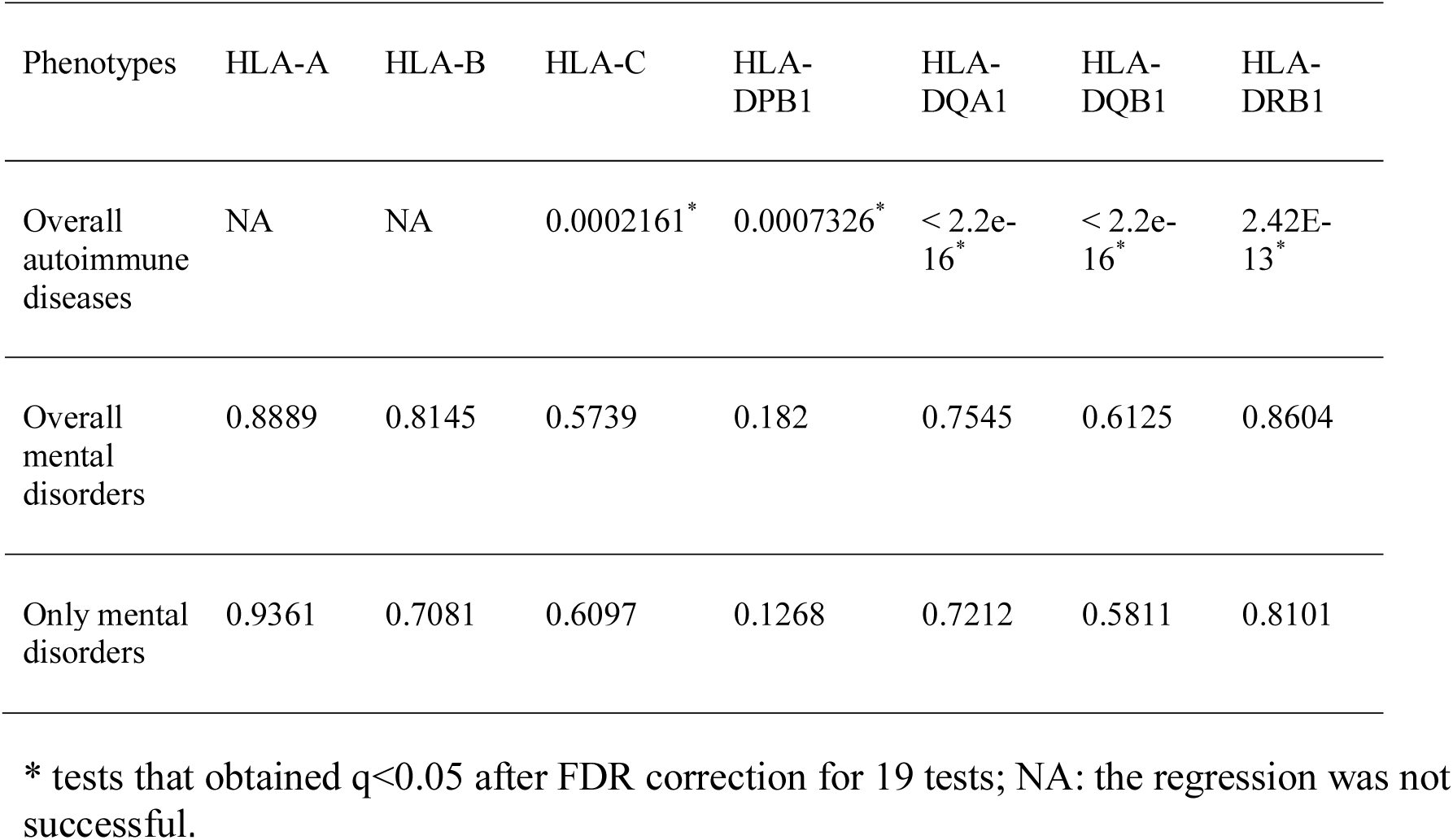
Association *P* values of HLA genes with autoimmune and mental disorders using gene-based tests

## Discussion

To our knowledge, this study is the largest prospective cohort sample systematically examining the overall epidemiological association between autoimmune and mental disorders. Compared to previous studies which focused on the associations between some specific diseases, we extended our analyses by including more people born between 1981 and 2012, investigating the bidirectional association between a variety of both autoimmune diseases and mental disorders, as well as the underlying genes explaining the associations. Our results replicated the overall comorbidity and bidirectional risk patterns of the two illness classes. Though GWAS for individual autoimmune diseases have been performed (**Table 3**), we have for the first time conducted a GWAS for overall autoimmune diseases, crossing seven specific diseases, and we observed that MHC region, especially HLA genes are genome-wide significantly associated with overall autoimmune diseases, but we did not find significant evidence for the connection of these genes with overall mental disorders, which were confirmed by our HLA allele association analysis.

Even though there is an overall positive epidemiological correlation between the two disease classes, when we examined the relationship between overall autoimmune diseases and specific mental disorders, or overall mental disorders and specific autoimmune diseases, the results vary. This is in line with previous reports, strongly indicating that the complex relationship between them should be cautiously addressed. The HLA region has been suggested to be the most plausible pathogenic interrelation between autoimmune and mental disorders. These analyses were however, restricted to specific disease associations, such as schizophrenia and rheumatoid arthritis (25). Our systematical analyses show that HLA genes were not the shared susceptibility loci with mental disorders in general, which has been replicated when we performed the HLA gene-based tests, demonstrating again the blurred and delicate associations between the two diseases. However, there could still be subgroups of the individuals with mental disorders but no autoimmune diseases diagnosis that have HLA associations and potentially undetected autoimmune diseases, due to the large heterogeneity of mental disorders and underlying pathophysiology.

Among the 20 HLA alleles from the five genes that were associated with overall autoimmune diseases, we observed that the components of the HLA class II genes, *HLA-DRB1, DQA1* and *DQB1* have the strongest associations, and all the alleles from each gene show either increased or decreased frequencies in patients with at least one autoimmune disease, with 13 in total increasing the risk of developing autoimmune diseases. It is also noteworthy that the top five alleles were DQB1*0302 (OR=1.77, 95%CI=1.60-1.95, *P*=3.16×10^-30^), DQA1*0301 (OR=1.74, 95%CI=1.56-1.94, *P*=1.23×10^-23^), DQB1*0602 (OR=0.57, 95%CI=0.50-0.65, *P*=3.06×10^-16^), DRB1*0401 (OR=1.70, 95%CI=1.48-1.94, *P*=1.28×10^-14^) and DRB1*1501 (OR=0.59, 95%CI=0.51-0.68, *P*=1.72×10^-12^), which were the frequent alleles detected in several autoimmune diseases, including type 1 diabetes, rheumatoid arthritis and multiple sclerosis (26). Specifically, in the case of diabetes, both DQB1*0302 and DQA1*0301 were associated with increased risk of type 1 diabetes (27); DQB1*0602 was reported to have a protective effect in type 1 diabetes (28, 29). Similarly, DRB1*0401 was found to increase risk of type 1 diabetes (30). DRB1*1501, as part of a larger haplotype, was found to be protective in type 1 diabetes (31). All these results are in line with the trends identified in our study. Conversely, a haplotype which includes both DQB1*0602 and DRB1*1501, both protective in our study, was found to increase risk of multiple sclerosis in a Swedish sample (32). This result attests to the complexity of the mechanism through which the HLA loci influence disease susceptibility and has been reported in previous studies of autoimmune diseases (33, 34). Thus, our study provides further evidence of the associations within HLA region with autoimmune diseases, and reports new associations with overall autoimmune diseases, increasing our understanding of the mechanisms behind disease pathogenesis.

Polygenic risk scores (PRSs) aggregate information from a large number of potentially causal genetic variants and hence is expected to be powerful for the risk prediction of polygenic traits, such as autoimmune and mental disorders. In our large, population-based study, we applied PRSs for autoimmune diseases to predict the disease status of mental disorders, and *vice versa*. Although PRSs were significantly associated with the respectively autoimmune or mental disorder phenotypes, our analysis did not show that autoimmune PRSs predict the individual risk of mental disorders, and mental PRSs can neither predict risk of autoimmune diseases. This finding is consistent with the previous findings by Euesden *et al.* (4) based on the 1958 British birth cohort study, who found that depression/rheumatoid arthritis genetic risk fails to predict rheumatoid arthritis /depression onset, although rheumatoid arthritis onset is associated with increased subsequent hazard of depression onset and *vice versa*. Thus, these analyses point to that even though PRSs capture a cumulative effect of multiple variants impacting autoimmune/mental disorders, they have the limited clinical utility for the disease onset prediction of mental/autoimmune diseases.

As the number of controls is 45 times of patient cases with overall autoimmune diseases in our cohort, to test whether our study results are sensitive to the extreme unbalanced size between cases and controls. For that, we randomly selected five controls per autoimmune case on the basis of various factors, including year of birth, gender and mental disorder status using propensity score matching method (35), and compared the OR and 95%CI values between full (i.e. 1,383 cases *versus* 62,656 controls) and sub samples (i.e. 1,383 cases *versus* 6,915 controls). We observed no statistical difference between the two results (**Figure S2**), which evidences the robustness of our outcome.

The strengths of our study include the clinically diagnosed indicators for both disease status, and the use of iPSYCH2012 cohort which recruited participants of homogeneous Danish descent with large sample size, thereby minimizing the risk of population stratification confounding factor. Though our sample is large for a single psychiatric cohort, we have low sample sizes for autoimmune cases. Because of this, our study has several limitations. Firstly, we were not able to estimate the heritability of overall autoimmune diseases, and the genetic correlation with overall mental disorders. Also, we were unable to investigate the specific relationship between the two diseases due to the low sample size of some specific autoimmune diseases. Moreover, we did not find significant association beyond the HLA region (chromosome 6 in the region between 26MB and 34MB). Larger studies are therefore needed to thoroughly investigate the genetic pleiotropy between the two diseases. Finally, our study focused exclusively on the genetic risk factors explaining the overall comorbidity, however, environmental exposures or an interaction of both should not be ignored. Cigarette smoking, for instance has been reported to be associated with both mental disorders, such as major depressive disorder (36), and also autoimmune diseases, such as rheumatoid arthritis (37). Therefore, future studies would be needed to better address the possibility of interactions with environment, and further improve understanding of disease pathobiology.

In conclusion, in our Danish population-based case-cohort sample (iPSYCH2012) including 1,383 patients with autoimmune diseases and 43,902 with mental illness, we replicated the finding that the two disease classes are often to be comorbid on the same patients. Furthermore, we for the first time systematically examined the overall bidirectional relationship between mental and autoimmune disorders. Later, our GWAS and HLA imputation analyses reveal the HLA genes are significantly associated with overall autoimmune diseases, but not with overall mental disorders.

## Contributors

MB conceived and designed the project. XL carried out the epidemiological, statistical genetics bioinformatics analyses, and wrote the first draft of the manuscript. RN performed the HLA association analysis. WT supervised the statistical analyses of the HLA study. SR and RA performed the imputation and QC of HLA alleles. VA, AS and AB performed the data cleaning and genetic variants imputation for iPSYCH2012 cohort. XL, RN and MB discussed and interpreted results. TW, PW, OM, DH, AB and MN are the principal investigators of the iPSYCH cohort. All authors contributed to the final manuscript.

## Acknowledgements

All personal information from the registers is anonymized when used for research purposes, and the project was approved by the Danish Data Protection Agency, hence according to Danish legislation; informed consent from participants was not required. This study was funded by The Lundbeck Foundation, Denmark (grant numbers R102-A9118, R268–2016–3925, and R155–2014–1724), the Independent Research Fund Denmark (grant number 7025–00078B), and the Mental Health Services Capital Region of Denmark, University of Copenhagen, Aarhus University and the university hospital in Aarhus. Genotyping of iPSYCH samples was supported by grants from the Lundbeck Foundation, the Stanley Foundation, the Simons Foundation (SFARI 311789), and NIMH (5U01MH094432–02). This research has been conducted using the Danish National Biobank resource supported by the Novo Nordisk Foundation. RA and SR were supported by the Novo Nordisk Foundation (NNF14CC0001). The iPSYCH data were stored and analyzed at the Computerome HPC Facility (http://www.computerome.dtu.dk/).

