## Supplementary File for "Genetic factors underlying the bidirectional relationship between autoimmune and mental disorders – findings from a Danish population-based study"

**Table of contents**

**Supplementary Figure 1:**	Hazard ratios of overall mental disorders with 95%CI	**2**

**Supplementary Figure 2:**	Comparison of effect sizes between full and sub samples.	**3**

**Supplementary Table 1:**	Complex associations between overall autoimmune and mental disorders	**3**

**Supplementary Table 2:**	Allele-specific tests for the significant genes	**4**

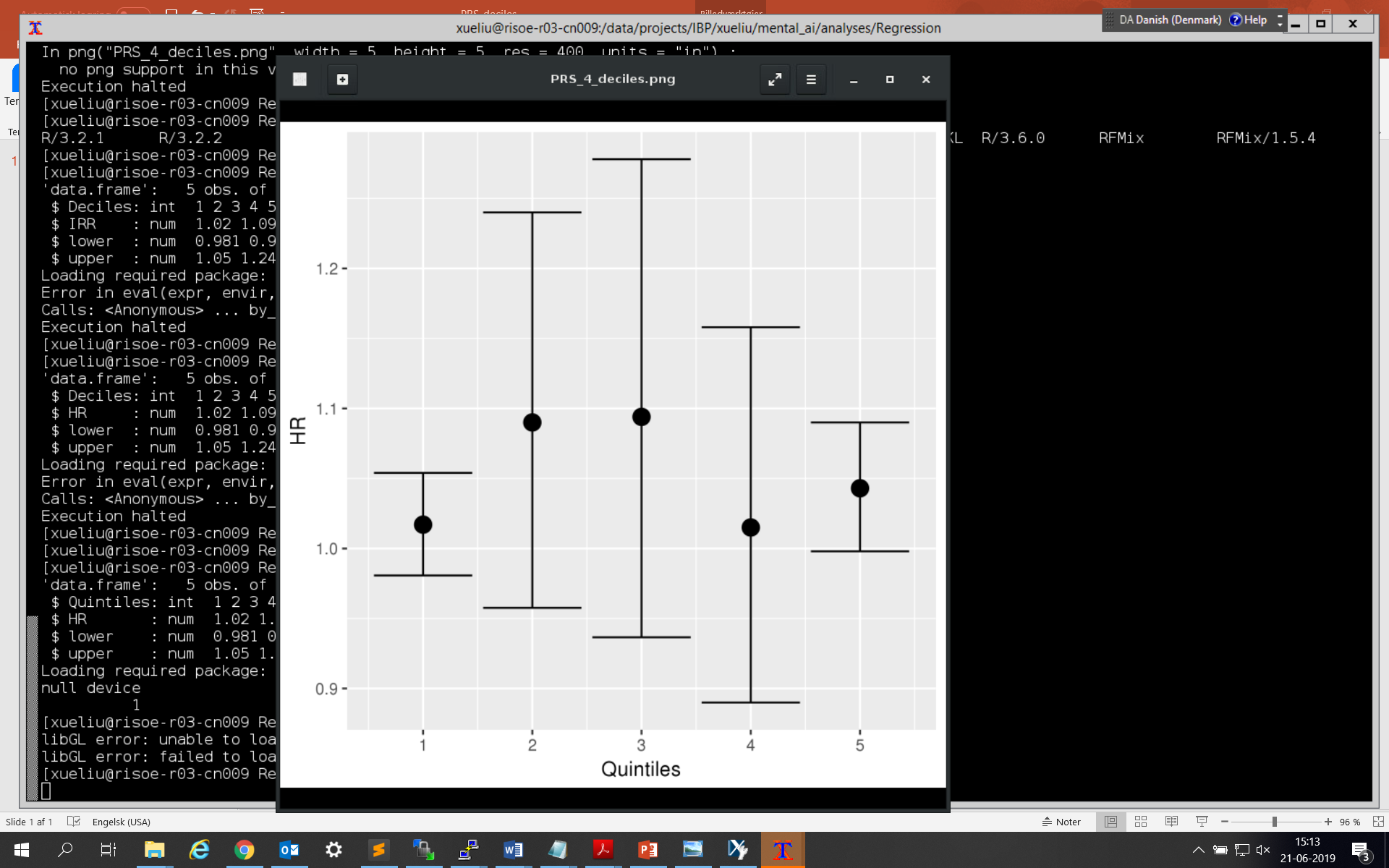

**Supplementary Figure 1**: Hazard ratios of overall mental disorders with 95%CI, associated with polygenic risk score quintiles for autoimmune diseases (adjusted for sex and four principal components).

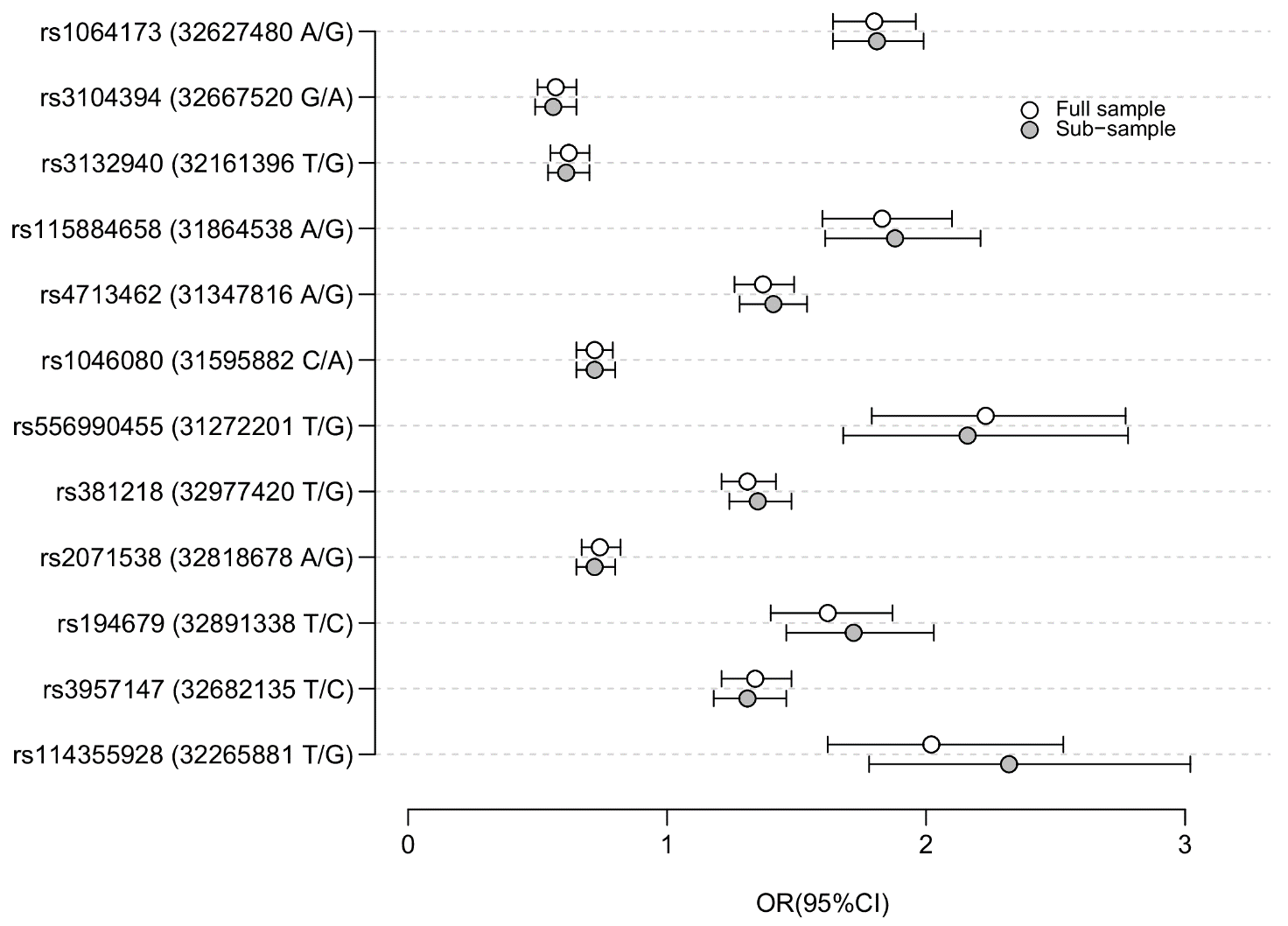

**Supplementary Figure 2.** Comparison of effect sizes between full and sub samples. Full sample indicates all the cases with overall autoimmune diseases and controls in the iPSYCH2012 cohort (i.e. 1,383 cases vs. 62,656 controls). Sub-sample includes all the cases and five controls per case (i.e. 1,383 cases vs. 6,915 controls). The y axis contains 12 independent genetic variants, positions and effect/other alleles are shown in brackets.

**Supplementary Table 1.** Complex associations between overall autoimmune and mental disorders

| Association testing* | HR | 95%CI | *P* (unadjusted) |
| --- | --- | --- | --- |
| Overall mental vs. Overall autoimmune disorders | 1.13 | 1.07,1.21 | 7.95e-05 |
| Overall autoimmune vs. Overall mental disorders | 1.27 | 1.16,1.39 | 8.77e-15 |
| Schizophrenia vs. Overall autoimmune disorders | 1.52 | 1.19,1.92 | 0.000797 |
| Overall autoimmune disorders vs. Schizophrenia | 2.64 | 2.03,3.44 | 3e-15 |
| Bipolar disorder vs. Overall autoimmune disorders | 1.07 | 0.73,1.55 | 0.737 |
| Overall autoimmune disorders vs. Bipolar disorder | 1.62 | 1.09,2.41 | 0.0104 |
| Affective disorders vs. Overall autoimmune disorders | 1.22 | 1.12,1.32 | 1.15e-05 |
| Overall autoimmune disorders vs. Affective disorders | 1.33 | 1.18,1.49 | 7.97e-10 |
| Autism spectrum disorders vs. Overall autoimmune disorders | 1.2 | 1.04,1.38 | 0.0118 |
| Overall autoimmune disorders vs. Autism spectrum disorders | 1.64 | 1.39,1.95 | 1.41e-14 |
| ADHD vs. Overall autoimmune disorders | 1.03 | 0.9,1.19 | 0.627 |
| Overall autoimmune disorders vs. ADHD | 1.9 | 1.61,2.24 | 0 |
| Anorexia vs. Overall autoimmune disorders | 1.37 | 1.12,1.67 | 0.000653 |
| Overall autoimmune disorders vs. Anorexia | 1.32 | 1.04,1.66 | 0.00282 |
| Overall mental disorders vs. CD | 1.03 | 0.89,1.2 | 0.645 |
| CD vs. Overall mental disorders | 0.97 | 0.78,1.19 | 0.578 |
| Overall mental disorders vs. UC | 0.86 | 0.75,0.99 | 0.0109 |
| UC vs. Overall mental disorders | 0.86 | 0.71,1.04 | 0.0111 |
| Overall mental disorders vs. CeD | 1.87 | 1.53,2.29 | 2.42e-07 |
| CeD vs. Overall mental disorders | 2.19 | 1.6,2.99 | 1.9e-09 |
| Overall mental disorders vs. RA | 1.09 | 0.88,1.35 | 0.37 |
| RA vs. Overall mental disorders | 1.07 | 0.78,1.47 | 0.423 |
| Overall mental disorders vs. PSOR | 1.09 | 0.9,1.32 | 0.336 |
| PSOR vs. Overall mental disorders | 1.12 | 0.85,1.48 | 0.239 |
| Overall mental disorders vs. T1D | 1.21 | 1.09,1.35 | 0.00173 |
| T1D vs. Overall mental disorders | 1.55 | 1.32,1.81 | 9.99e-16 |
| Overall mental disorders vs. JIA | 1.42 | 1.21,1.65 | 0.000235 |
| JIA vs. Overall mental disorders | 1.88 | 1.5,2.37 | 3.21e-12 |

*: we evaluated the bidirectional associations between two diseases. When we evaluated A *vs.* B, then in this model we estimated the effect of B on the outcome A. Likewise, when we evaluated B *vs.* A, then we estimated effect of A on the outcome B. CD: Crohn’s disease. UC: ulcerative colitis. CeD: celiac diseases, RA: rheumatoid arthritis. PSOR: psoriasis. T1D: type 1 diabetes. JIA: juvenile idiopathic arthritis.

**Supplementary Table 2.** Allele-specific tests for the significant genes

| Alleles | OR(95%CI) | *P* | q-value* |
| --- | --- | --- | --- |
| DQB1*0302 | 1.77(1.6-1.95) | 3.16E-30 | 1.93E-28 |
| DQA1*0301 | 1.74(1.56-1.94) | 1.23E-23 | 3.76E-22 |
| DQB1*0602 | 0.57(0.5-0.65) | 3.06E-16 | 6.23E-15 |
| DRB1*0401 | 1.7(1.48-1.94) | 1.28E-14 | 1.95E-13 |
| DRB1*1501 | 0.59(0.51-0.68) | 1.72E-12 | 2.10E-11 |
| DQA1*0102 | 0.73(0.66-0.81) | 3.99E-09 | 4.06E-08 |
| DQA1*0401 | 1.53(1.27-1.85) | 8.60E-06 | 7.51E-05 |
| DQB1*0402 | 1.48(1.24-1.77) | 1.90E-05 | 0.00014 |
| C*0304 | 1.24(1.11-1.38) | 0.00012 | 0.00080 |
| DPB1*0301 | 1.27(1.12-1.45) | 0.00031 | 0.0019 |
| DQA1*0501 | 1.28(1.11-1.46) | 0.00043 | 0.0024 |
| DPB1*0402 | 0.76(0.65-0.88) | 0.00049 | 0.0024 |
| DQB1*0501 | 0.78(0.68-0.9) | 0.00052 | 0.0024 |
| DQB1*0201 | 1.26(1.1-1.43) | 0.00054 | 0.0024 |
| C*0401 | 0.75(0.64-0.89) | 0.00069 | 0.0028 |
| DRB1*0801 | 1.46(1.17-1.83) | 0.00084 | 0.0029 |
| C*0202 | 1.31(1.12-1.54) | 0.00086 | 0.0029 |
| DRB1*0301 | 1.3(1.11-1.52) | 0.00087 | 0.0029 |
| DRB1*0405 | 2.92(1.43-5.95) | 0.0033 | 0.01 |
| C*0702 | 0.85(0.76-0.95) | 0.0033 | 0.01 |

*, multiple testing corrected by false discovery rate method.
